# Guidance on filtering DNA methylation microarray probes by detection p-values

**DOI:** 10.1101/245217

**Authors:** Jonathan A. Heiss, Allan C. Just

## Abstract

DNA methylation microarrays are popular for epigenome-wide association studies, but spurious outlier values complicate downstream analysis. It was demonstrated previously that commonly used detection p-value cut-offs were insufficient and led to many calls on the Y-chromosome probes in females. We extend these observations by assessing 2,578 samples from 18 studies run on the 450K chip as well as outliers across technical replicates. We provide comprehensive guidance and software for filtering methylation microarrays as an essential step to reduce outliers.

## Introduction

The Illumina Infinium HumanMethylation450 (450K) and the more recent Infinium MethylationEPIC (850K) arrays are two widely popular platforms for epigenome-wide association studies. Before beginning with downstream analyses, comprehensive quality control (QC) should be conducted to identify problematic samples. But samples passing QC still contain outliers, e.g. when targeted loci are present in low quantity due to amplification artifacts, or are mutated and no longer match their probe sequence. Affected probes feature mostly background noise and should be excluded. This decision is based on detection p-values, a concept explained below. Suggested cut-offs in the literature span several magnitudes from 0.05 to 1e-16. Lehne et al. systematically evaluated cut-offs based on the idea that probes targeting the Y-chromosome should be detected among males but not females [1]. We provide updated guidance with a corrected implementation across multiple datasets.

## Methods

Treating DNA with bisulfite converts epigenetic modifications into distinct base sequences. In combination with subsequent whole-genome amplification, the ratio of methylated/unmethylated CpG sites is translated into differences in abundance of these distinct sequences which hybridize to complementary probes on the microarray and are linked with fluorescent dyes. By comparing fluorescence intensities of the probes targeting the methylated (*M*) and unmethylated (*U*) variant of a CpG site, its methylation level in the DNA input can be inferred. In case of a completely unmethylated CpG site, the *U* probe features a very high intensity, whereas the *M* intensity is low, or vice versa in case of a completely methylated CpG site. Thus, the total intensity *T*=*U*+*M* is — to a certain degree — independent of the methylation level itself. High intensities usually indicate good signal-to-noise ratio and such probes are hence deemed *detected*, whereas low intensities indicate *undetected* probes. Where to draw the line is based on the concept of detection p-values: the parameters of a background distribution *B* are estimated based on probes thought to feature mainly background noise. If the observed value of *T* is unlikely to be generated by *B* (associated p-value is below significance level) a probe is considered detected, otherwise undetected. The previous recommendation by Lehne et al. is based on *minfi* v1.2.0 [2] which misspecified the parameters of B (adding standard deviations instead of variances).

We explored two approaches to estimate *B* using either negative control probes (NEG), specifically designed not to match the human genome, or so-called out-of-band (OOB) intensities, i.e. those observed in the usually unused color channel [3]. Whereas former approach uses raw intensities as currently implemented in *minfi*, latter approach involves a dye-bias correction step (using the RELIC method [4]) before detection p-values are computed. Robust estimators of location and spread were used. 18 public 450K datasets comprising 2,667 samples representing a wide range of tissues were downloaded. 2,578 samples passed our comprehensive QC including screening for sex-mismatches. Samples were grouped by sex (m/f=1,220/1,358) and the number of detected Y-chromosome probes was counted for a range of p-value cut-offs.

Code to reproduce analyses is provided in the supplement and a function to calculate detection p-values can be found in the *ewastools* package (github.com/hhhh5/ewastools).

## Results

Fig. 1 shows the median (2.5th and 97.5th percentiles) number out of the 416 Y-chromosome probes that were classified as detected grouped by sex. When estimating *B* using NEG evaluated cut-offs ranged from 1e0 to 1e-80, and from 1 to 0.001 for the higher OOB intensities as they consequently resulted in higher p-values (to achieve similar results for NEG as for the combination OOB/0.05 a cut-off around 1e-117 was necessary, see Fig. 2).

**Figure 1:**
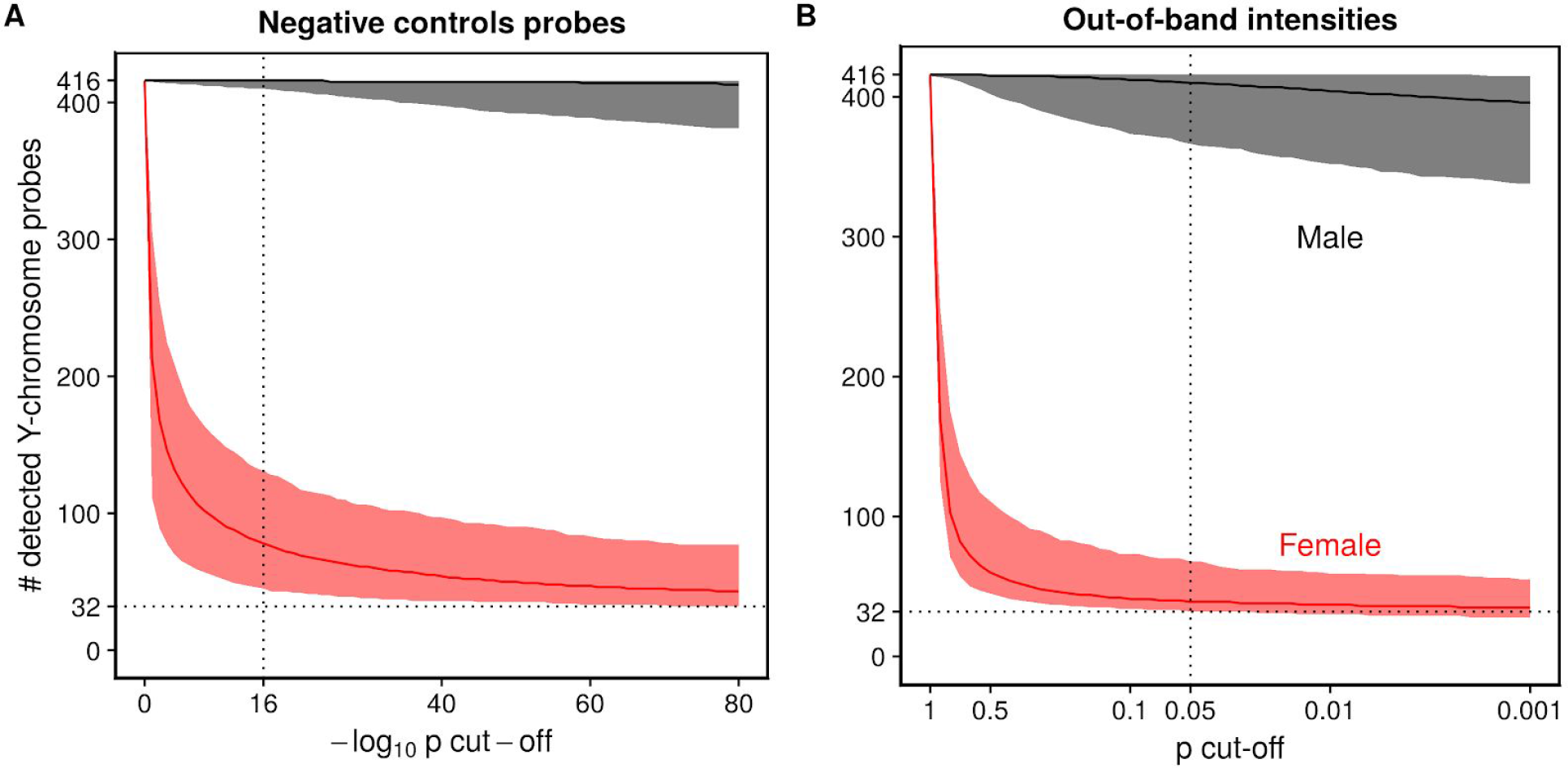
Median number and 95% interval of detected Y-chromosome probes among male and female samples for various detection p-value cut-offs: (**A**) using negative controls probes to estimate background distribution; (**B)** using out-of-band intensities.

**Figure 2:**
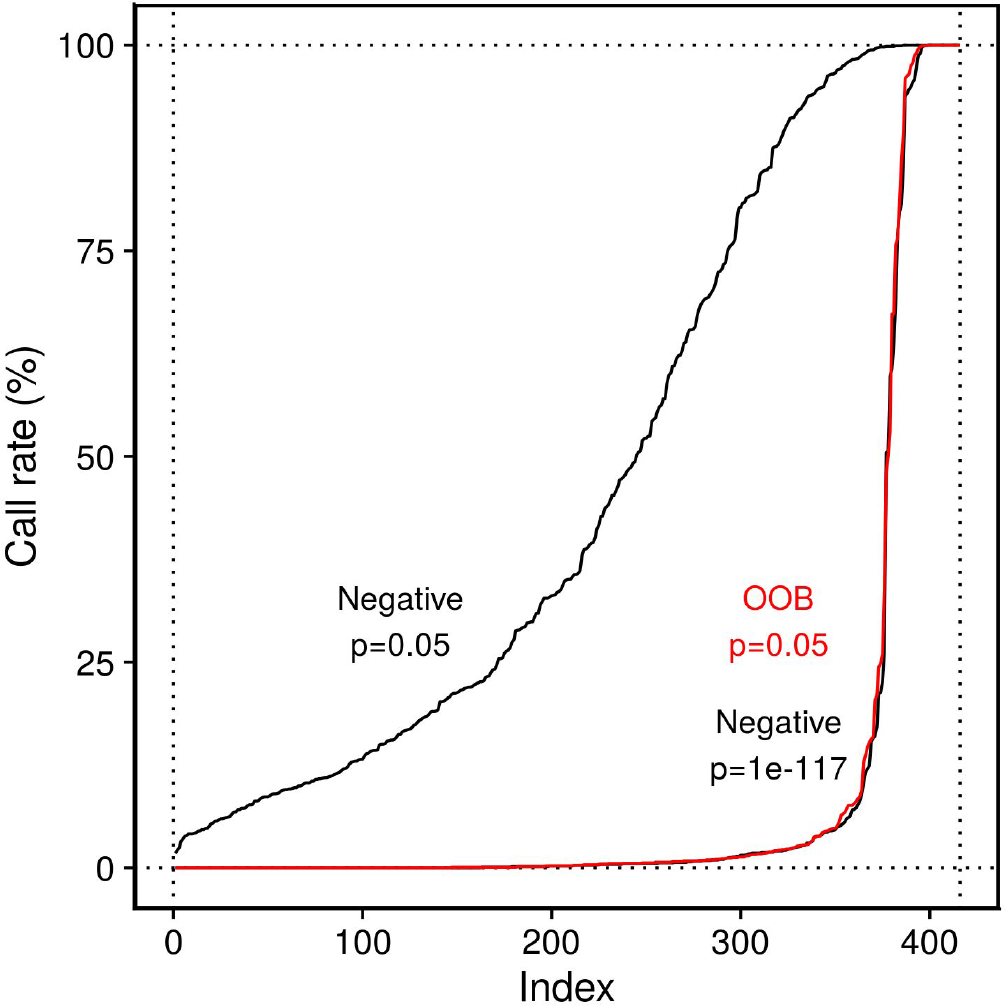
Detection rates of Y-chromosome probes among the 1,358 female samples. For the common threshold of 0.05 and a background distribution estimated from negative control probes, not a single probe had a detection rate of zero. For more stringent criteria there is a almost clear-cut separation between undetected and detected (probably cross-reactive with autosomal locprobes. The order of probes is not identical between curves.

Even for the most stringent cut-offs considered here the number of detected probes among females was non-zero, but when calculating the detection rate among females for each probe (Fig. 2) there was an almost clear-cut separation between always undetected and detected (probably cross-reactive with autosomal loci) probes.

Using NEG and a cut-off of 0.05 resulted in a median of 64 undetected probes per sample, not counting Y-chromosome probes. Lowering the cut-off to 1e-16 increased this number to 332 probes. Switching to OOB and a cut-off of 0.05 resulted in a median of 3,780 undetected probes per sample, approximately 0.8% of the data.

A subset of 6 index samples (whole blood) and their technical replicates was used to assess the absolute difference in methylation levels for paired measurements. The median difference was 1.3 percentage points (pp) among the pairs with probes detected in both index sample and replicate, and 7.7pp (NEG/0.05), 6.2pp (NEG/1e-16) and 5.0pp (OOB/0.05) for all other pairs, respectively. 3,733 pairs had a difference >20pp, of which 283 (8%) (NEG/0.05), 1058 (28%) (NEG/1e-16) and 2,576 (69%) (OOB/0.05) were classified as undetected, respectively.

## Conclusions

While both negative control probes and OOB intensities are utilized in background subtraction/correction, they show very distinct distributions. OOB intensities might reflect more realistic levels of background noise resulting from cross-hybridization as many probes possess some sequence similarity with off-target loci resulting in off-target binding. In contrast, negative control probes are designed not to match the human genome (their probe sequences are proprietary) and consequently feature lower intensities. We recommend to use OOB to estimate the background distribution after dye-bias correction with a detection p-value cut-off of 0.05, as this combination helped to exclude the majority of major outliers (>20pp) between technical replicates. Most undetected probes showed smaller differences and removing such data points may seem overly conservative, however, with the exception of a few exposures such as smoking, effect sizes of associations in whole blood are often very small and removing unreliable data points may strengthen statistical power. Examples include BMI [5], diabetes [6], or age, all of which were found and replicated to be associated with methylation changes of a few percentage points.

